# Evolution of contribution timing in public goods games

**DOI:** 10.1101/2020.01.09.900670

**Authors:** Bryce Morsky, Marco Smolla, Erol Akçay

## Abstract

Life history strategies are a crucial aspect of life, which are complicated in group-living species, where payoffs additionally depend on others’ behaviours. Previous theoretical models of public good games have generally focused on the amounts individuals contribute to the public good. Yet a much less studied strategic aspect of public good games, the timing of contributions, can also have dramatic consequences for individual and collective performance. Here, we develop game theoretical models to explore how the timing of contributions evolves. We show how contributing rapidly is not necessarily optimal, since delayers can act as “cheats,” avoiding contributing while reaping the benefits of the public good. However, delaying too long can put the delayers at a disadvantage as they can miss out on the benefits. These effects lead to bistability in a single group, and spatial diversity among multiple interacting groups.

## 1 Introduction

In all biological systems, timing plays a fundamental role, from molecular-level circadian rhythms controlling gene expression, to large-scale migratory movement and breeding timing. The right timing allows proteins to be ready by the time they are needed (e.g. digestive enzymes), to arrive at foraging or breeding sites at an advantageous time, or to exhibit a specific phenotype at the right time. Exogenous stimuli, such as temperature (Sims et al., 2001), are often involved in coordinating behaviours. Moreover, where large numbers of individuals coordinate behaviours, timing can also create issues, such as, crowding and competition for limited resources (Skoglund et al., 2012). In fact, modelling approaches suggest that density-dependence can lead to a timing diversification to avoid competition (Metcalf et al., 2015).

In group living animals, timing issues can lead to specific social dilemmas, where the interests of the group are in conflict with the interests of the individual. An example is when individuals contribute to a public good which benefits the group as a whole, such as, predator vigilance, provisioning of offspring, and sustainable resource use. Sometimes these benefits are only achieved if sufficient amounts of time or resources have been contributed to the public good: meerkat (*Suricata suricatta*) sentinel behaviour (Bednekoff, 2015; Clutton-Brock et al., 2001), mitigating climate change (Milinski et al., 2008), and herd immunity (Fine et al., 2011) are examples. If sufficient individuals contribute to the public good, everyone will be better off. However, given that contributing generally comes at a personal cost, such as resources used in provisioning, time spent guarding, or putting oneself in danger, an individual will be better off if it does not contribute while others do. This undermines cooperation and ultimately leads to a collapse of contributions. How social and biological systems can prevent the collapse of such public goods has been the focus of much of economic and game theoretic literature (Cadsby and Maynes, 1999; Pacheco et al., 2008; Tavoni et al., 2011).

Previous work generally assumes that participants in a public goods game differ in their behavioural strategies, which determine the magnitude of their contribution at a fixed point in time (Pacheco et al., 2008; Wang et al., 2009). An individual’s payoff, or fitness, is typically the benefit that the individual incurs from the public good minus the cost of contributing. However, another important aspect in public goods games, the timing of contributions, has received relatively little attention (with some exceptions Chakra and Traulsen, 2012; Hilbe et al., 2013; Chakra et al., 2018). For example, in pharaoh ant colonies, *Monomorium pharaonis*, many potentially unrelated queens live together and cooperate (Schmidt et al., 2011). Queens continuously produce queen, king, and worker eggs. However, workers curate these eggs and will only rear worker eggs until the colony grows and the egg to worker ratio falls below a certain threshold (Warner et al., 2018). Here, workers represent a public good to the queens as they rear the offspring, but the benefit from the public good in terms of reproductive offspring is only realized when the colony is big enough. Queens that delay expending energy into worker production will have more reproductive offspring (queens) and therefore higher fitness compared to those that contribute early on. Strikingly, the timing of reproductive effort can be influenced by *Wolbachia* endosymbionts (Singh and Linksvayer, 2019). Similarly, in the greater ani, several pairs of this communally breeding bird will protect a shared nest, incubate eggs, and provide food to chicks. While generally cooperative, females will roll eggs out of the nest until they have started to lay their own eggs and so reduce competition for their offspring (Riehl and Jara, 2009). Thus, females that lay their eggs later will avoid having their eggs ejected (Riehl, 2010). Cooperation and public goods games are also important in the biology of microorganisms (West et al., 2006; Tarnita, 2017). In particular, the behaviour of altruistic sacrifice where some cells forgo reproduction to the benefit of others such as in the formation of fruiting bodies under food deprivation in the slime mold *Dictyostelium discoideum* (Strassmann et al., 2000). Timing of contribution into the fruiting body can play a role in whether the microorganism is sacrificed into forming the stalk or forms the spores. In an environment of collaborators, fast production of a stalk would be the social optimum. Yet, delaying may permit a cell to be a member of the head of the stalk. Timing plays a role here: wait too long, and the spores will be formed or even spread; too early, and you contribute to the stalk.

In these examples, even if everyone contributes the same amount, there is a conflict over the timing of contributions. If delaying contributions means more resources available for private benefits or competition once a public good is produced, individuals will have an incentive to delay their contributions. On the other hand, delayed contributions means that the group as a whole will have to wait longer until the public good is produced, and so the benefits from the public good are realised more slowly. In some cases, delaying contribution can also run afoul of time constraints, such as seasonal changes or predation risk, where fitness will be zero if contributions have been delayed for too long.

This conflict over the timing of contributions to public good can have knock-on effects on life-history decisions. Life-history strategies commonly describe the trade-off between investing resources, such as energy and time, into growth or reproduction. Whatever is not invested into growth is what is left for reproduction and vice versa. However, where fitness depends on the timely production of a public good, fitness may also depend on the timing of contributions relative to others. Therefore, in social organisms when components of fitness depend on the production of public goods, life history schedules might be affected by the conflict over timing of contribution.

It is therefore important to understand how conflicts over the timing of contribution to public goods play out in evolutionary games. However, previous work generally assumed that individuals always invest at the same time but may vary in the amounts they contribute. To study the effect of contribution-timing, we develop a threshold public goods model where members of a group expend effort (or contribute resources) towards a common good. Once, a specific total contribution level is reached, such as the construction of a nest, the group enters a different non-cooperative state, where individuals expend their effort into competition with one another over the common resource. We define an individual’s behavioural strategy as the timing of its contribution. That is, all individuals can contribute the same amount in principle, but vary in their schedule of when to make the contributions. This may mean that depending on others’ schedules an individual might end up contributing more or less than others until the public good is produced.

We use this model to explore under which conditions we would expect contributions to be delayed, and to investigate whether delayers can co-exist with non-delayers. For a single group, we find bistablity between the two strategies in the conflict over timing. We then extend this model to a spatial, multi-group setting, to test whether groups of delayers and non-delayers can co-exist. We find that the bistability of single group dynamics lead to spatial patterns of stable monomorphic clusters of delayers and non-delayers. Our results provide the first evolutionary game theoretic account for how conflict over timing of contributions to a public goods game plays out in well-mixed and spatially structured populations.

## 2 Methods

We begin with a model of a single group of players and a single public good. This model is a discrete dynamical system where contributions are made in continuous time (first stage) whereas payoffs are earned at discrete events once the accumulation of a specific amount of resources has been reached (second stage). In the second part, we use reaction-diffusion equations to study evolutionary group dynamics for multiple groups and multiple public goods games.

### 2.1 Single group and public good model

Consider a group of players contributing to a public good, such as group vigilance or cooperative breeding. We assume that each player has the same amount of resources to contribute over their life-times and that contributions follows a hump-shaped ‘energy schedule’ over time, *f* (*t*). That is, *f* (*t*) starts out small, peaks partway through, and then declines as the individual senesces. This reflects the biologically realistic assumption that the performance of behaviors or phenotypes typically will need to increase from a low or zero baseline, and will decline later in life. The actual shape of the performance schedule does not qualitatively affect our results, so long as the curves are unimodal. As we are interested in contribution timing, individuals play one of two strategies: a ‘delayed schedule’ or ‘non-delayed schedule.’ The delayed schedule is shifted along the time axis by an amount *δ*. Thus *f* (*t* − *δ*) and *f* (*t*) are the energy schedules for delayers and non-delayers, respectively.

In our model (Fig. 1) all efforts flow towards the public good until the threshold amount of resources is met at which point the group enters the second stage of the game, which is a non-cooperate scramble for resources (e.g. opportunities to reproduce). We assume that the scramble happens relatively quickly, so that it depends only on the performance at the time the threshold is reached.

**Figure 1.**
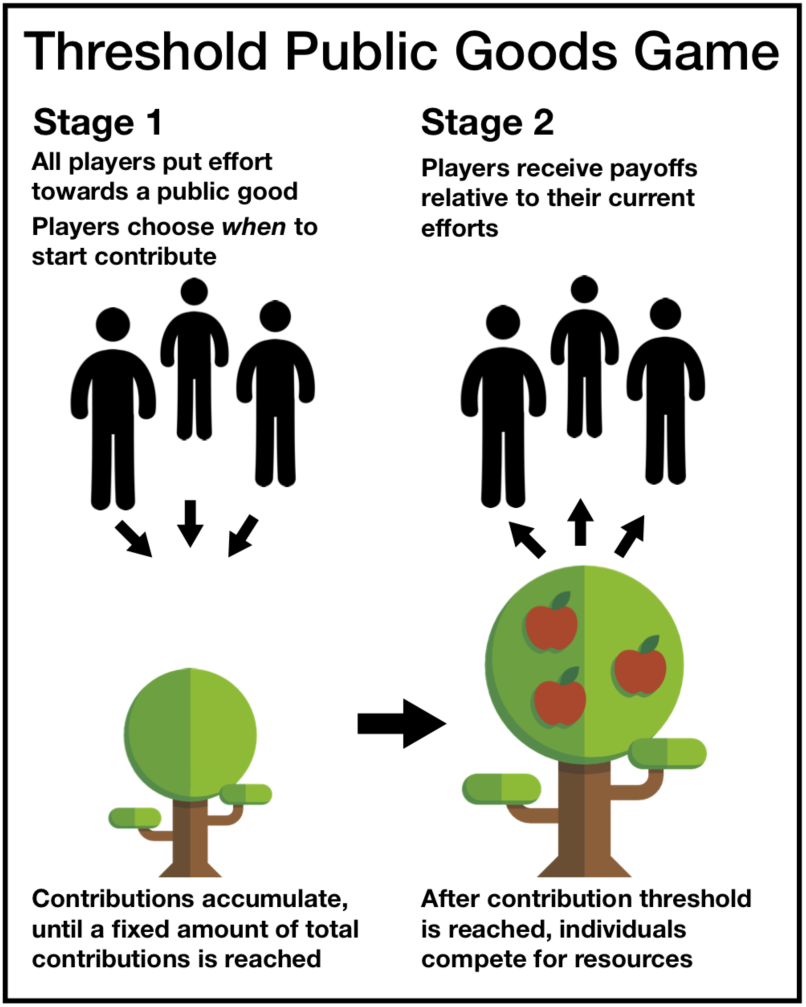
Schematic of the threshold public goods game.

To model the evolution of this process, we employ a discrete dynamical system that tracks the proportion of delayers, *p*_*n*_, and the time, *θ*_*n*_, at which the condition is met for generation *n*. The condition is met when a total contribution level *C* is reached by the population, i.e.

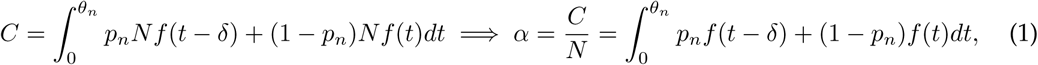

where *N* is the number of individuals in the group and *α* the total per capita contribution. We then solve Eq. 1 for *θ*_*n*_. We calculate the proportion of the delaying players at the next generation *p*_*n*+1_ as

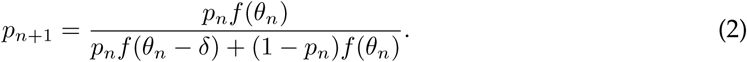

We also considered a fixed time threshold model, where *θ* is a fixed parameter. See Supplementary Information A for an analysis of this model.

### 2.2 Multiple groups and public goods model

Spatial dynamics can have important effects upon life-history traits, such as in host-pathogen systems (Lion and Gandon, 2015). To explore the impact of delaying, we extend the previous model by letting multiple groups play their own dynamic public goods game in a spatial setting. We assume that these groups are spread out in a square two-dimensional space with periodic boundaries. Each point in space has a carrying capacity, and players may randomly migrate to adjacent areas.

To model the growth dynamics as a result of social interactions together with migration, we use a reaction diffusion model that extends the model in Sec. 2.1 with the addition of growth rates and a carrying capacity. Reaction diffusion models are well studied models of spatial ecology (Cantrell and Cosner, 2004). For simplicity, we assume carrying capacity to be constant across space. We track the number of delayers, *n*_1_, and non-delayers, *n*_2_, at each point in space. Additionally, we take the time until payoffs are earned, *θ*, in each group into account. If a group has a lower *θ* relative to another group, it will reproduce sooner and so grow more quickly.

Let us first consider the continuous time dynamics at a single location before we extend the model to a spatial version. We use a two-type Lotka-Volterra system that takes both the growth rates and carrying capacity, *K*, into account. This approximates the single group model at a single point, which we discussed above. The change of *n*_1_ and *n*_2_ over time are given by:

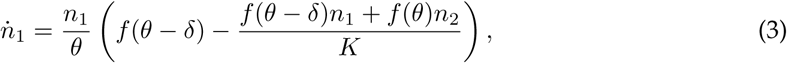

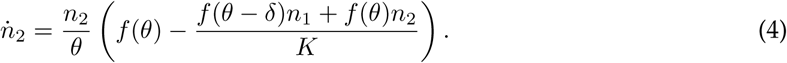

Note that dividing by *θ* takes the generation time into account, i.e. the time until payoffs are earned, thus inducing a cost to delaying contribution into the public good in one group relative to another group. The smaller *θ*, the faster the population will grow. *θ* is constrained by Eq. 1, making this a differential algebraic equation (DAE).

In the next step, we extend this continuous time approximation to multiple locations. Each location has an *x, y* coordinate. Thus, the number of delayers and number of non-delayers at (*x, y*) and time *t* is given by *n*_1_(*x, y, t*) and *n*_2_(*x, y, t*) respectively. To simulate movement of players between patches, we add a diffusion term to Eqs. 3 and 4, which yields the following reaction diffusion equations:

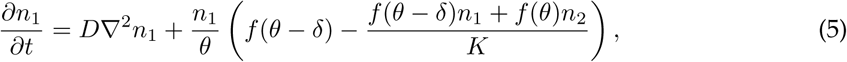

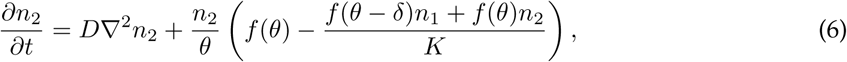

where *D* is the diffusion constant, ∇ is the spatial gradient, and ∇^2^ is the Laplace operator. Finally, *θ* is determined by local patch conditions via Eq. 1, yielding a partial differential-algebraic equation (PDAE). We solved this system numerically. See Supplementary Information B for details.

## 3 Results

### 3.1 The single group model

For the single group model, we find that the groups will either be all delaying or all non-delaying at equilibrium (*p*\*, *θ*\*). The exception is where *f* (*θ*\* − *δ*) = *f* (*θ*\*). This is an edge-case where the dynamics devolve into *p*_*n*+1_ = *p*_*n*_, where any mix of both types is an equilibrium. Fig. 2 portrays the stability of the monomorphic equilibria for low, intermediate, and high values of *α*. When *α* is low, the times *θ* at which the thresholds are met, for delayers and non-delayers (Fig. 2A,D respectively), are short. As such, the all non-delaying equilibrium is stable, while the all delaying one is not. For intermediate levels of *α*, however, we observe bistability as both equilibria may be stable (Fig. 2B,E). For high *α*, only a monomorphic population of delayers is stable (Fig. 2C,F). See Supplementary Information C.1 for a mathematical analysis of these equilbria.

**Figure 2.**
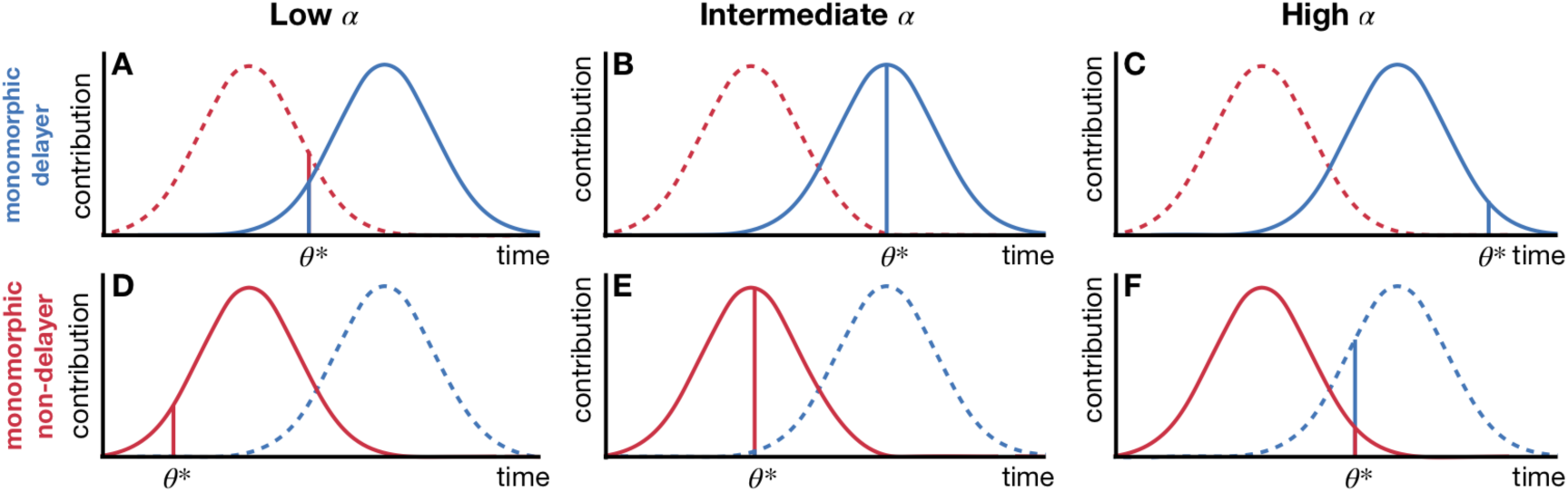
A graphical representation of the stability of the monomorphic equilibria for varying *α* (columns) and the delayed and non-delayed strategies (rows). Vertical bars indicate the relative contribution level of delayers (blue) and non-delayers (red) at *θ*\* when the contribution threshold *α* is reached. All delaying is stable where *α* is high (C), whereas the all non-delaying is stable where *α* is low (D). For intermediate values of *α*, we have bistability as both strategies are stable at this point (B,E). The invading mutant strategies are shown for each panel as dashed curves with the respective colour.

Although the stability of the monomorphic equilibria depends on *α*, where a polymorphic equilibrium exists in a single group, it is always unstable (see Supplementary Information C.1). The dependence of the two strategies on *α* is shown in Fig. 3 as a bifurcation diagram. For low *α*, there is only one equilibrium, monomorphic non-delaying. As *α* increases, the monomorphic delaying equilibrium becomes stable and an unstable polymorphic equilibrium emerges. Further increasing *α* destabilizes the monomophic non-delaying equilibrium. We lose the polymorphic equilibrium and are left with the stable monomorpic delaying equilibrium. The range of *α* where bistability obtains is larger when *δ* is larger.

**Figure 3.**
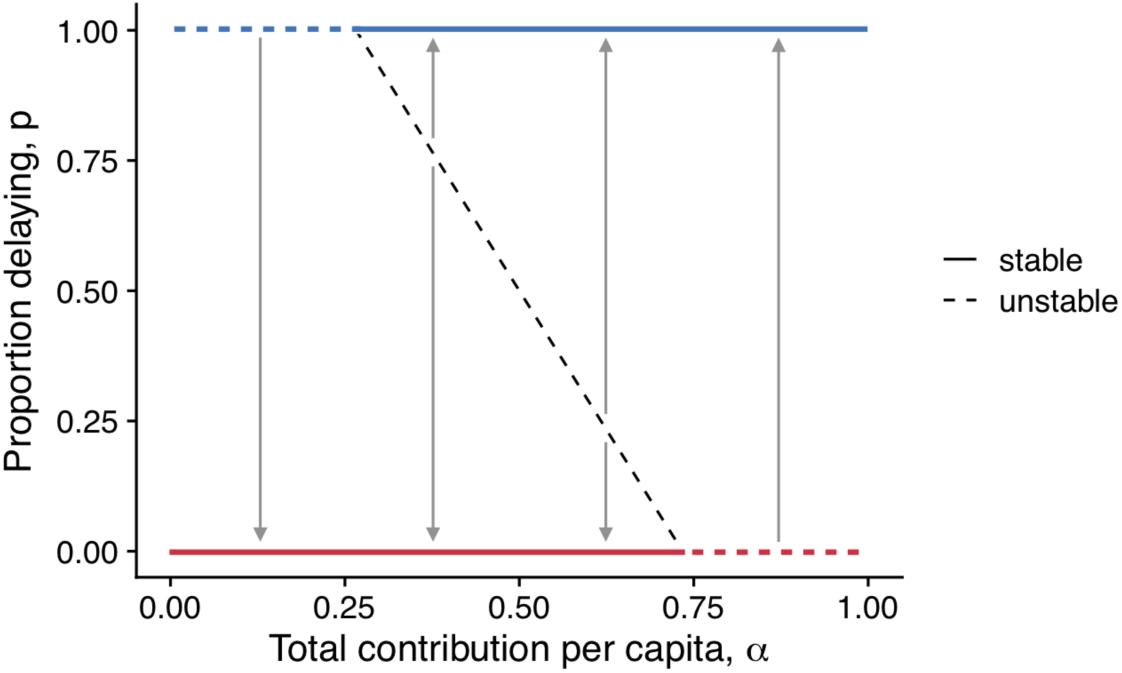
The bifurcation diagram shows that when *α* is low, non-delayers (pink) dominate, while for high *α*, delayers (blue) dominate, whereas for intermediate values of *α* the system is bistable. Shown are results where the performance function, *f* (*t*), is the probability distribution function of the Normal distribution 𝒩(*μ* = 10,*σ* = 2), and *δ* = 1.25*μ* (25% delay). Solid lines indicate stable states, dashed lines indicate unstable states.

We analysed the adaptive dynamics of this system, and found that for a low contribution threshold *α* where *f* (*θ*\*) is less than the peak value, expediting the schedule will evolve. If, however, the contribution threshold is high, *f* (*θ*\*) is greater than the peak value, and so further contribution delaying will evolve. In these cases. evolution will only halt once some boundary effect is reached, i.e. delaying or expediting cannot be done indefinitely (see Supplementary Information C.2 for an analysis).

### 3.2 The multi-group reaction-diffusion model

The Lotka-Volterra model for a single group (Eqs. 3 and 4) behaves similarly to the single group model, i.e. bistability is contingent on *α* (see Supplementary Information D). To analyse the spatial extension, we numerically solved the fixed-contribution model for a variety of initial conditions and parameter values. The energy functions *f* (*t*) and *f* (*t* − *δ*) are normal probability distribution functions (PDF). For non-delayers, the mean is *μ* = 10 and the standard deviation is *σ* = 2. We consider delayers with several different delays, *δ* = 10%, 30%, and 50% of *μ* (i.e. *f* (*t* − *δ*) is a normal PDF with mean *μ*(1 + *δ*)). We also consider *α* = 0.4, 0.5, and 0.6 and diffusion constant *D* = 0.01. For greater or lower *α*, the population fixes to one or the other type. Initial conditions were *N* = 5, and the proportion of delayers drawn from an *L*×*L* Gaussian field (see Supplementary Information B for details). We explored the effects of spatially uncorrelated initial conditions using a multivariate Beta distribution with parameters (0.5, 0.5), i.e. a U-shaped distribution on [0, 1], see Supplementary Information E.2.

In Fig. 4, we show heatmaps of the proportion of delayers across space at time *t* = 1000 for different values of *α* and *δ*. Note the coexistence and patterns that occur. For *α* outside of the bistable region, we do not observe such patterns as the population evolve to a spatial monomorphism (results not shown here). The delay parameter *δ* has an impact on how sharp the boundaries are between clusters of the two types are. When the delay is low, the boundaries are fuzzier than when it is higher. This is because increasing the delay increases local bistability resulting in patches of (nearly) all delayers or all non-delayers. This effect amplifies the formation of clusters, which can then be maintained. As observed in Fig. 4 panels F and I, while there are no polymorphisms for *α* = 0.6 and *δ* = 30%, there are if the delay is increased to *δ* = 50%.

**Figure 4.**
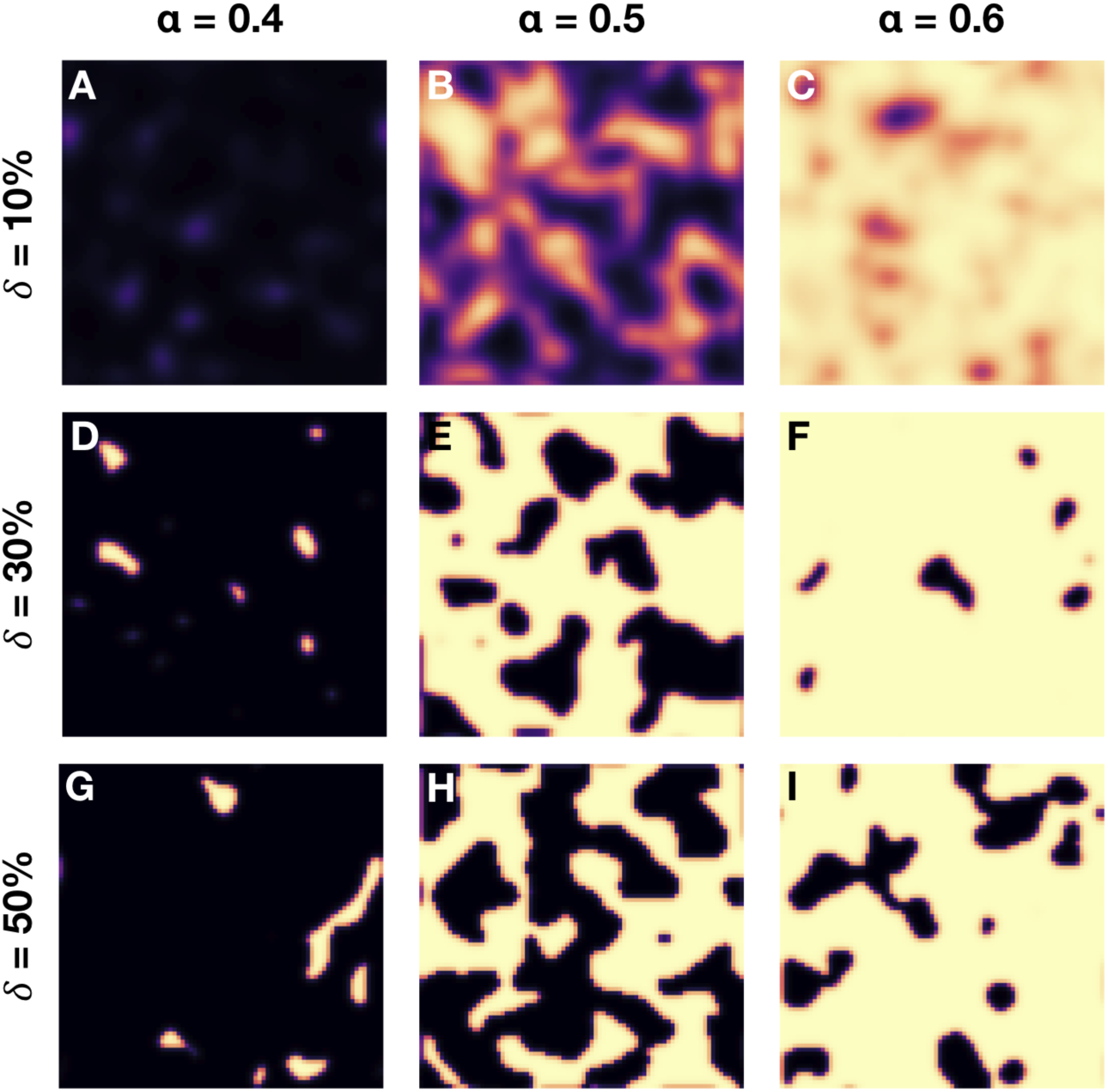
Delayers and non-delayers co-exist for a range of parameters. Shown are the spatial polymorphisms of delayers (yellow) and non-delayers (black) for the multi-group, multi public goods game model for low, intermediate, and high *α* (columns), and where delayers delay contributions with *δ* ∈ {10%, 30%, 50%}. These snapshots are taken at time *t* = 1000.

Fig. 5 depicts the time series for various parameter combinations. The time to reach equilibrium is faster as *α* is high or low. Convergence is slowest in the middle of the bistable region at *α* = 0.5. Unlike the discrete dynamical system in the single group model, non-delayers provide a benefit to their group with a higher growth rate via the division by *θ* (which is lower the higher the proportion of non-delayers). Thus, groups with more non-delayers will grow faster than groups with fewer, and thereby spread more quickly throughout space. However, as the carrying capacity is reached, this effect abates and the local dynamics dominate (see Supplementary Information E.1 for a time series of spatial mean of *N*). This process leads to the fluctuations in the time series most evident for longer delays. Early on, growth dominates and non-delayers spread. However, as the carrying capacity is reached, the local dynamics can force the groups into more delayers.

**Figure 5.**
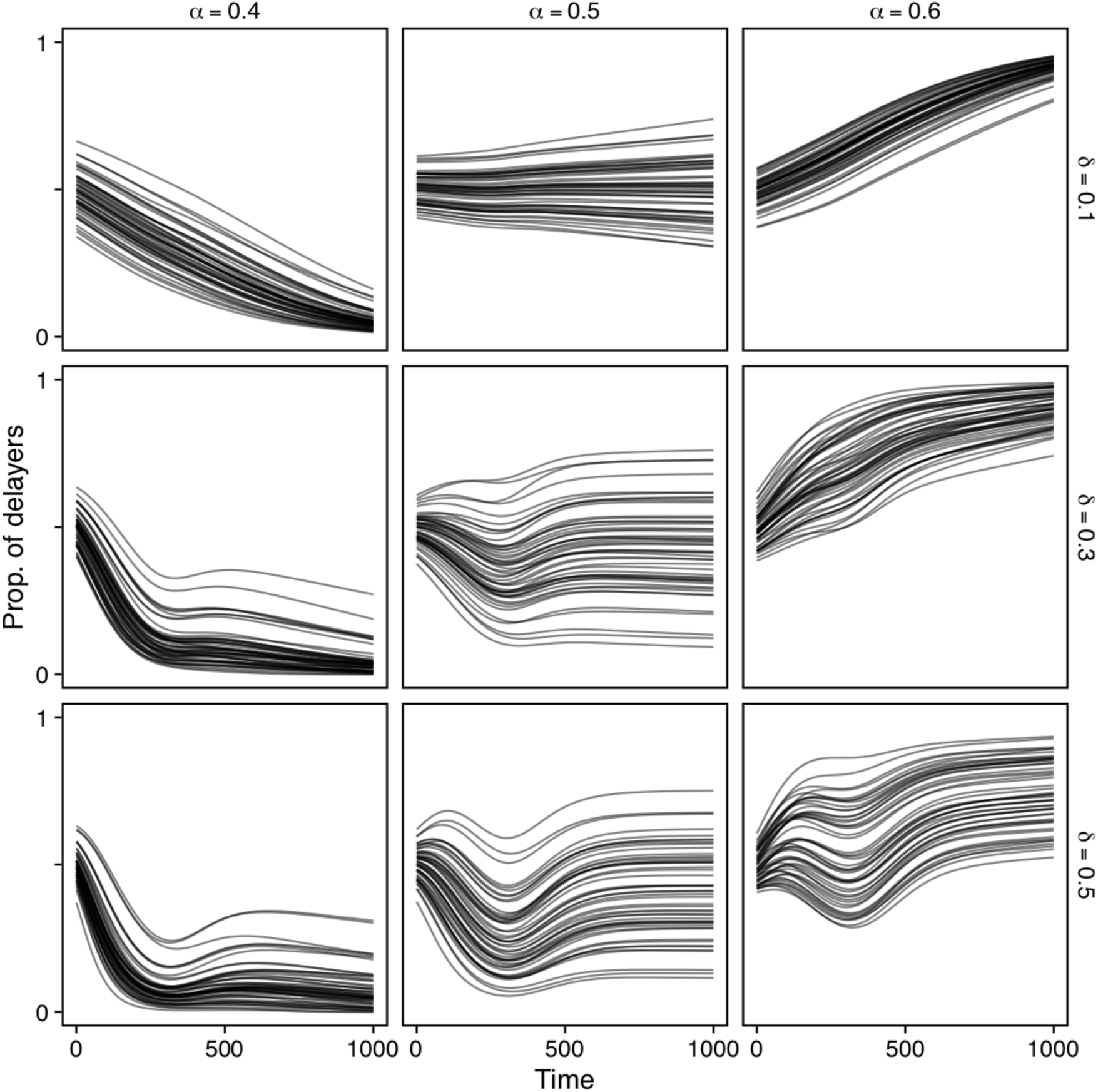
50 time series of the spatial average of *p* for low, intermediate, and high *α* (columns), and where delayers delay contributions with *δ* ∈ {10%, 30%, 50%}. *D* = 0.01. The initial proportion of delayers is a normalized Gaussian random field.

The time series in Fig. 5 also show sensitivity to initial conditions for intermediate *α*. At *α* = 0.5, we see that the population can evolve to be predominantly delayers or non-delayers, depending on the initial conditions. For initial conditions close to equal proportions of each type, we observe the most coexistence.

## 4 Discussion

Timing is important in biology, especially in social settings where not just what individuals do but also when they do it can have important payoff consequences. We investigated these consequences in a threshold public goods game, where players contribute efforts towards a public good for which they will compete at some point. Our model sheds light on how the tension between individual- and group-level incentives for early vs. delayed contribution plays out at the group and population level. Intuitively, contributing early should be beneficial as it allows the public good to be realised earlier, which should help a non-delaying strategy to spread. However, a player that is delaying contributions can free-ride on others’ contributions. While this sounds like a classic social dilemma, it comes with a twist. For intermediate levels of required contribution, it is best to be in sync with the majority of players as they will primarily determine the groups’ dynamics. This happens because individuals’ payoffs are determined by their performance when the public good is realised. Thus, each individual has an incentive to be at their peak performance when the threshold is reached, which means that when most group members contribute early, it is best to also contribute early, and conversely if most group members delay contributions, it is best to also delay. When we consider the continuous evolution of contribution timing we find a similar result: either contributing as early as possible or delaying as much as possible is selected for, depending on where the contribution threshold lies relative to the peak of the energy schedule. This positive frequency dependence and bistability are in contrast to the instantaneous threshold public goods games where the incentives typically give rise to negative frequency dependence. When groups exist in a spatially distributed population, this local bistability can result in a pattern of patchy coexistence of delayers and non-delayers.

A few previous studies (Chakra and Traulsen, 2012; Hilbe et al., 2013; Chakra et al., 2018) looked at the role of timing of contributions in collective-risk games such as the Volunteer’s Dilemma (Diekmann, 1985). In these models, players observe each others’ behaviour over several turns before benefits are accrued and contributions are contingent on the behaviour of others (Chakra and Traulsen, 2012; Hilbe et al., 2013). However, because the payoffs are a function of the total amount contributed plus the amount that is withheld, not contributing always increases an individual’s payoff. In the models of Chakra and Traulsen (2012) and Hilbe et al. (2013), the payoffs do not directly depend on the timing of contributions, as payoffs are realised at the end of the interaction and depend only on the total contributions. But, timing might affect the responses of individuals to each other. They find that strategies that delay contributions are selected for. In contrast, Chakra et al. (2018) consider a model inspired by abatement actions to avert climate change, where in each round there is a risk of incurring loss that players can abate by contributing to the public good. They show that such ongoing risks can incentivize early contributions, because current contributions mean higher expected endowments in later rounds. Our model highlights a different strategic issue that can arise with timing when contributions have to follow a performance curve that needs to be ramped up first. We find that similar to other public goods games, delayers can act as cheaters, avoiding contributing to a public good while reaping its benefit. However, if the public good is realised early, then delayers can “miss out” on capitalising on the public good, and thereby lose to non-delayers who contributed more.

Our model is also related mathematically to the truncation selection dynamics (Morsky and Bauch, 2016, 2019). In these models, variation in social interactions lead to hump-shaped payoff distributions for each strategic type. Fitness is only realised over some payoff threshold, and is the same for all individuals over the threshold. The main difference between these dynamics and our model is that under truncation selection, the total fitness of a strategic type is determined by integrating the entire remainder of the payoff distribution above the threshold. In our model, this corresponds to letting the competitive phase of the game last for a long time. Thus, truncation selection of Morsky and Bauch (2016) and our model can be seen as extremes of a family of models where the competitive phase is taken to last for some duration after the threshold is met: in our model this duration is infinitesimal, while Morsky and Bauch (2016) takes it as infinite. The shorter this competition duration is, the stronger the positive frequency dependence we uncover in our model will be.

Our spatial model allows us to study the dynamics of between group competition, where the outcomes depend on the initial distribution of delayers and non-delayers. Notably, we observe two distinct dynamical regimes in our simulations, which we start below the carrying capacity. In the first regime, groups grow to saturate the landscape, where the non-delayers have an advantage because of their faster growth rate, as indicated by the initial dip (or slow increase) in the frequency of delayers in Fig. 5. However, as populations reach carrying capacity, migration starts to be more important, and what happens at this stage depends on the clumpiness of the distribution of delayers and non-delayers: large clusters of both delayers and non-delayers can at least temporarily resist invasion by the other type, which results in a persistent polymorphism across space. These results have implications for maintenance of life-history variation: recent work on steelhead trout showed that variation in life history speed (spawning timing and parity) was maintained on a single river system by negative frequency dependence (Christie et al., 2018). Our results suggest that when life-history is a function of social interactions and threshold public goods, variation in speed can be maintained across space through positive frequency dependence.

An interesting application of our model are Pharaoh ants infected with the endosymbiotic bacteria, *Wolbachia*. Pharaoh ant queens, regardless of infection status, will contribute workers to the colony as a public good, forgoing reproduction until a threshold of workers to queens is reached. *Wolbachia* can accelerate this life cycle of the queens (Singh and Linksvayer, 2019). One would expect then that *Wolbachia* prevalence should increase. However, in the wild, both monomorphic infected and monomorphic non-infected colonies coexist in proximity to one another, unlike non-social insects such as mosquitoes, where spatial coexistence is not predicted from models (Barton and Turelli, 2011). The bistability and spatial patterns of our model can explain this observation.

Furthermore, timing of reproduction is frequently a crucial component of reproductive isolation and potential for speciation (Quinn et al., 2000; Hendry and Day, 2005). Our bistability result suggests a new mechanism in which timing differences in social behaviours can lead to reproductive isolation. Two sub-populations that evolved different timing strategies (e.g., in response to different thresholds in different environments) might not be able to interbreed even after they come in contact and experience the same environment.

The models that we have explored here can be expanded in a variety of ways such as unrealised public goods, and observability of contributions. In our model, individuals cannot change their performance trajectory nor can they adapt to others’ behaviours. An extension to this model could incorporate feedback wherein players may observe and then adjust their life-histories. Further, consider the case where the contributions into the public good dissipate over time, i.e. the common pool into which players contribute is leaky. In this case, delaying imposes a further cost to the group and erodes past contributions, and thereby the threshold might never be reached. Relaxing our other assumptions may have interesting results. For example, we assumed that the benefit from the public good is relative to the growth rates at the threshold in the single group model and the growth rates and time in the multi-group model. Further, we assumed that the carrying capacity was fixed i.e. it was neither a function of the total amount nor the timing of the contributions. Relaxing these assumptions would mean that group size and the contribution amount would be important. Different strategies could be favoured at different population sizes, and it would be important to explore the evolution of the threshold contribution level, *C*. Such a model could be further expanded by incorporating a mortality risk over time. There is much further work to explore the role of timing in evolutionary and ecological games with implications for biodiversity, speciation, and cooperation.

In conclusion, our work illustrates the interesting interactions between timing, life histories, and public goods games. We have shown that the impact of delaying life history is non-trivial. High contribution games foster delaying, while low ones foster advancing life histories. We have opened up new dimensions for both public goods games, and for life history strategies. More broadly, it is important to study the constraints that tie individuals together, and their impact on evolution and ecology.

## Acknowledgements

The authors would like to thank Rohini Singh and Tim Linksvayer for helpful discussions that motivated this study, as well as Jeremy Van Cleve for his technical support.

## Funding

This research was supported by the Army Research Office (W911NF-17-1-0017) and the University of Pennsylvania.

## Author contributions

All authors contributed to the conception of the study. B.M. did the mathematical analysis and led the development of the simulation code. B.M. and M.S. analysed the data and wrote the first draft, and all authors contributed to the final manuscript.

## Competing interests

The authors declare that they have no competing interests.

## Notes

https://github.com/erolakcay/contributiontiming

