## Supplementary Information for "Evolution of contribution timing in public goods games"

### Supplementary Information for
### Evolution of contribution timing in public goods games

Bryce Morsky [*], Marco Smolla [†], and Erol Akçay [‡]

Department of Biology, University of Pennsylvania, Philadelphia, PA, USA

January 9, 2020

#### A Analysis of a fixed-time condition model

Consider a fixed-time threshold model, i.e. $\theta$ is fixed. In addition to the monomorphic equilibria $p^* = 0, 1$, there are polymorphic equilibria when $f(\theta - \delta) = f(\theta)$. This case, however, is trivial, since the dynamics devolve to $p_{n+1} = p_n$. So, considering the case where $f(\theta - \delta) \neq f(\theta)$, we have

$$\left|\frac{d\phi}{dp_n}\right| = \left|\frac{f(\theta-\delta)f(\theta)}{(\bar{p}f(\theta-\delta) + (1-p^*)f(\theta))^2}\right| \implies \left|\frac{d\phi}{dp_n}\right|_{p_n=0} = \left|\frac{f(\theta-\delta)}{f(\theta)}\right|, \quad \left|\frac{d\phi}{dp_n}\right|_{p_n=1} = \left|\frac{f(\theta)}{f(\theta-\delta)}\right|, \quad (1)$$

where

$$\phi(p_n) = p_{n+1} = \frac{p_n f(\theta_n)}{p_n f(\theta_n - \delta) + (1-p_n) f(\theta_n)} \quad (2)$$

is the discrete map. Therefore, if $f(\theta - \delta) < f(\theta)$, then $p^* = 0$ is stable. If $f(\theta - \delta) > f(\theta)$, then $p^* = 1$ is stable.

With respect to the spatial version of this model, one of the two strategies will have a higher growth rate (except if they are exactly equal) and will therefore dominate. Thus, the spatial model will be qualitatively the same as the non-spatial one.

#### B Numerical methods

To numerically solve the reaction-diffusion model, we use the method of lines. We discretize space into an $L \times L$ lattice with $n_{1,j,k}$ and $n_{2,j,k}$ the number of delayers and non-delayers at position $(j, k)$, and consider periodic boundaries. The finite difference scheme for the diffusion operator is

[*]
[†]
[‡]

$$\nabla^2 n_{i,j,k} \approx \left(\frac{n_{i,j+1,k} - 2n_{i,j,k} + n_{i,j-1,k}}{h^2} + \frac{n_{i,j,k+1} - 2n_{i,j,k} + n_{i,j,k-1}}{h^2}\right) \qquad (3)$$
$$= \frac{1}{h^2}(n_{i,j+1,k} + n_{i,j-1,k} + n_{i,j,k+1} + n_{i,j,k-1} - 4n_{i,j,k}).$$

To numerically solve this system, we use Julia's DifferentialEquations package (Rackauckas and Nie, 2017) with solver Rodas4P, a Rosenbrock method (Hairer and Wanner, 1996). Our code is available at `https://github.com/erolakcay/contributiontiming`.

### C Analysis of the fixed contribution threshold model

#### C.1 Stability analysis

Here we discuss the equilibria and the stability of our discrete time model presented in the main text where the public good is produced when a fixed threshold contribution is reached (the time of which depends on the group composition). In addition to the monomorphic equilibria $p^* = 0, 1$, there are polymorphic equilibria when $f(\theta^* - \delta) = f(\theta^*)$. To determine stability, let us first define the maps $p_{n+1} = \phi_1(p_n, \theta_n)$ (Eq. 2) and $\theta_n = \phi_2(p_n)$. Noting that $\theta_n$ is determined via the condition

$$\alpha = \int_0^{\theta_n} p_n f(t - \delta) + (1 - p_n) f(t) dt \qquad (4)$$

as a function of $p_n$, we may simplify the model into a one dimensional map, $p_{n+1} = \phi_1(p_n, \phi_2(p_n))$. To check for stability, we differentiate with respect to $p_n$ and find

$$\left|\frac{d\phi_1}{dp_n}\right|_{p_n=1} = \left|\frac{\partial \phi_1}{\partial p_n} + \frac{\partial \phi_1}{\partial \phi_2}\frac{d\phi_2}{dp_n}\right|_{p_n=1} \qquad (5)$$

where (recalling that $\theta^* = \phi_2(p^*)$)

$$\left.\frac{\partial \phi_1}{\partial p_n}\right|_{p_n=p^*} = \frac{f(\theta^* - \delta) f(\theta^*)}{(p^* f(\theta^* - \delta) + (1 - p^*) f(\theta^*))^2}, \qquad (6)$$
$$\left.\frac{\partial \phi_1}{\partial \phi_2}\right|_{p_n=p^*} = \frac{p^*(1 - p^*)(f'(\theta^* - \delta) f(\theta^*) - f'(\theta^*) f(\theta^* - \delta))}{(p^* f(\theta^* - \delta) + (1 - p^*) f(\theta^*))^2}. \qquad (7)$$

To find $d\phi_2(p_n)/dp_n$, differentiate Eq. 4 with respect to $p_n$ and rearrange, i.e.

$$\alpha = \int_0^{\theta_n} p_n f(t - \delta) + (1 - p_n) f(t) dt$$
$$\implies 0 = \int_0^{\theta_n} f(t - \delta) - f(t) dt + \frac{d\phi_2}{dp_n}(p_n f(\theta_n - \delta) + (1 - p_n) f(\theta_n)),$$
$$\implies \left.\frac{d\phi_2}{dp_n}\right|_{p_n=p^*} = \frac{\int_0^{\theta^*} f(t) - f(t - \delta) dt}{p^* f(\theta^* - \delta) + (1 - p_n) f(\theta^*)}, \qquad (8)$$

Checking stability for the monomorphic states gives us

$$\left|\frac{d\phi_1}{dp_n}\right|_{p_n=0} = \left|\frac{f(\theta^* - \delta)}{f(\theta^*)}\right| < 1, \quad \left|\frac{d\phi_1}{dp_n}\right|_{p_n=1} = \left|\frac{f(\theta^*)}{f(\theta^* - \delta)}\right| < 1. \tag{9}$$

If $\alpha$ is sufficiently high, then $f(\theta^* - \delta) > f(\theta^*)$. Therefore, the monomorphic delayer state is stable. The monomorphic non-delayer state can be stable if $\alpha$ is sufficiently low, since $f(\theta^* - \delta) < f(\theta^*)$. Thus, we observe contrasting bifurcations with respect to $\alpha$; increasing (decreasing) $\alpha$ causes instability for $p^* = 1$ ($p^* = 0$). The bifurcations occur at $f(\theta^* - \delta) = f(\theta^*)$. Moving onto the polymorphic state, where $f(\theta^* - \delta) = f(\theta^*)$, it is unstable for

$$\left|\frac{d\phi_1}{dp_n}\right|_{p_n=p^*} = \left|1 + \frac{p^*(1-p^*)(f'(\theta^* - \delta) - f'(\theta^*))(\int_0^{\theta^*} f(t) - f(t-\delta)dt)}{f(\theta^*)^2}\right| \geq 1 \tag{10}$$

$$\iff \frac{p^*(1-p^*)(f'(\theta^* - \delta) - f'(\theta^*))(\int_0^{\theta^*} f(t) - f(t-\delta)dt)}{f(\theta^*)^2} > 0. \tag{11}$$

Now, note that the denominator is positive, and note that $\int_0^{\theta^*} f(t) - f(t-\delta)dt > 0$, due to the delay. Further, $f(\theta^* - \delta) = f(\theta^*)$ implies that $f'(\theta^* - \delta) > 0$ and $f'(\theta^*) < 0$. Thus, the polymorphic state is unstable.

#### C.2 Evolutionary invasion analysis

Let us now consider the evolution of delaying within an adaptive dynamics framework. Rare mutations occur in $\delta$ such that the population has time to reach equilibrium before each new mutation. We check the invasibility of a mutant by differentiating the relative growth rates of the mutant to resident at the resident's equilibrium time $\theta^*_{\text{res}}$ (i.e. the eigenvalue Eq. 9) with respect to $\delta$ and evaluating at $\delta = 0$,

$$\frac{\partial}{\partial\delta}\left(\frac{f(\theta^*_{\text{res}} - \delta)}{f(\theta^*_{\text{res}})}\right)\Bigg|_{\delta=0} = -\frac{f'(\theta^*_{\text{res}})}{f(\theta^*_{\text{res}})}, \quad \frac{\partial}{\partial\delta}\left(\frac{f(\theta^*_{\text{res}} + \delta)}{f(\theta^*_{\text{res}})}\right)\Bigg|_{\delta=0} = \frac{f'(\theta^*_{\text{res}})}{f(\theta^*_{\text{res}})}, \tag{12}$$

for a relatively delaying and expediting mutant, respectively. A mutant can invade the equilibrium if the corresponding equation is positive. Thus, at $p^* = 0$, a mutant that expedites its schedule relative to the resident can invade if $f'(\theta^*_{\text{res}}) < 0$, i.e. $\theta^*_{\text{res}} > \theta^\dagger_{\text{res}}$, where $\theta^\dagger_{\text{res}}$ is the time at which the resident's schedule is at its peak. A mutant that delays relative to the resident can invade if $f'(\theta^*_{\text{res}}) > 0$, i.e. $\theta^*_{\text{res}} < \theta^\dagger_{\text{res}}$. If $\theta^*_{\text{res}} = \theta^\dagger_{\text{res}}$, then $f'(\theta^*_{\text{res}}) = 0$. Analysing the second derivative gives us

$$\frac{\partial^2}{\partial\delta^2}\left(\frac{f(\theta^*_{\text{res}} - \delta)}{f(\theta^*_{\text{res}})}\right)\Bigg|_{\delta=0} = \frac{f''(\theta^*\theta^\dagger_{\text{res}})}{f(\theta^*\theta^\dagger_{\text{res}})} < 0, \quad \frac{\partial^2}{\partial\delta^2}\left(\frac{f(\theta^*_{\text{res}} + \delta)}{f(\theta^*_{\text{res}})}\right)\Bigg|_{\delta=0} = \frac{f''(\theta^\dagger_{\text{res}})}{f(\theta^\dagger_{\text{res}})} < 0, \tag{13}$$

which means that it is an evolutionary stable point.

Note that a mutation that is too large may not be able to invade. Assume that the schedule is monotonically increasing until it peaks and monotonically decreasing thereafter. Then, for some $\theta^*_{\text{res}} > \theta^\dagger_{\text{res}}$, there exists a $\tilde{\theta} < \theta^\dagger_{\text{res}}$ such that $f(\theta^*_{\text{res}} - \delta) > f(\theta^*_{\text{res}})$ for $\delta > \theta^*_{\text{res}} - \tilde{\theta}$, and can thereby invade. A similar such situation occurs for $\theta^*_{\text{res}} < \theta^\dagger_{\text{res}}$.

Now, in our examples, we set the schedules to be normal distributions. Thus, expediting the schedule will reduce the total amount of energy over the individual's lifetime, and increase $\theta^*_{\text{res}}$. Therefore, if $\theta^*_{\text{res}} < \theta^\dagger_{\text{res}}$, mutants can invade until $\theta^*_{\text{res}} = \theta^\dagger_{\text{res}}$, at which point evolution halts. This effect does not necessarily occur in the other direction. Though delaying will decrease $\theta^*_{\text{res}}$, $\theta^\dagger_{\text{res}}$. However, a reasonable constraint is that one cannot delay indefinitely, and therefore beyond some time all energy is zero. In this case, $\theta^*_{\text{res}}$ will decrease until it reaches the peak, at which point evolution halts. Note that if we use a different schedule such that the total energy for an individual remains constant regardless of shifts in timing (i.e. shifts move the peak, but $f(0-\delta)$ is fixed for all $\delta$), then for $\alpha$ such that $\theta^*_{\text{res}} < \theta^\dagger_{\text{res}}$, the schedule approaches the delta function at $t = 0$. And, for $\alpha$ such that $\theta^*_{\text{res}} > \theta^\dagger_{\text{res}}$, the schedule approaches the delta function at $t = \infty$.

### D Analysis of the Lotka-Volterra DAE

#### D.1 Stability analysis

Here we will analyze our Lotka-Volterra model, which is a differential-algebraic equation (DAE) (Beardmore and Song, 1998). The model features a carrying capacity $K$, interspecific competition, and an algebraic constraint. Further, we show that this system exhibits bistability, like the discrete system.

The Lotka-Volterra DAE equations are

$$\begin{pmatrix} \dot{n}_1 \\ \dot{n}_2 \\ 0 \end{pmatrix} = \Phi = \begin{pmatrix} \frac{n_1}{\theta}\left(f(\theta-\delta) - \frac{(f(\theta-\delta)n_1 + f(\theta)n_2)}{K}\right) \\ \frac{n_2}{\theta}\left(f(\theta) - \frac{(f(\theta-\delta)n_1 + f(\theta)n_2)}{K}\right) \\ \int_0^\theta n_1 f(t-\delta) + n_2 f(t)dt - \alpha(n_1+n_2) \end{pmatrix}, \tag{14}$$

where $n_1$ and $n_2$ are the delayer and non-delayers, respectively. Following the derivation in Crow et al. (1970), the change in the proportion of delayers in a Lotka-Volterra equation is

$$\dot{p} = p(f(\theta-\delta) - pf(\theta-\delta) - (1-p)f(\theta)). \tag{15}$$

Now, scaling by $1/(p_t f(\theta-\delta) + (1-p_t)f(\theta)) > 0$, we have

$$\dot{p} \approx \frac{p_t(f(\theta-\delta) - p_t f(\theta-\delta) - (1-p_t)f(\theta))}{p_t f(\theta-\delta) + (1-p_t)f(\theta)} = p_{n+1} - p_n \tag{16}$$

$$\implies p_{n+1} = \frac{p_n f(\theta-\delta)}{p_n f(\theta-\delta) + (1-p_n)f(\theta)}. \tag{17}$$

To consider stability, we must first find the index of the DAE (Campbell and Gear, 1995). Differentiating the algebraic equation with respect to time and rearraranging with respect to $\dot{\theta}$, we find

$$\frac{d\phi_3}{dt} = \dot{n}_1\left(\int_0^\theta f(t-\delta)dt - \alpha\right) + \dot{n}_2\left(\int_0^\theta f(t)dt - \alpha\right) + \dot{\theta}(n_1 f(\theta-\delta) + n_2 f(\theta)) = 0 \tag{18}$$

$$\implies \dot{\theta} = \frac{\dot{n}_1\left(\alpha - \int_0^\theta f(t-\delta)dt\right) + \dot{n}_2\left(\alpha - \int_0^\theta f(t)dt\right)}{n_1 f(\theta-\delta) + n_2 f(\theta)}. \tag{19}$$

If $n_1 f(\theta-\delta)+n_2 f(\theta)\neq 0$, we have an index one DAE.

To check stability for the equilibia, we linearize $\Phi$, taking the Jacobian of $\Phi$ at an equilibrium, $J(n_1^*, n_2^*, \theta^*)$ (Riaza, 2004). Letting $M$ be the mass matrix, we then find the eigenvalues of the matrix pencil

$$\{M, J(n_1^*, n_2^*, \theta^*)\} = \left\{ \begin{pmatrix} 1 & 0 & 0 \\ 0 & 1 & 0 \\ 0 & 0 & 0 \end{pmatrix}, \left. \begin{pmatrix} \frac{\partial \phi_1}{\partial n_1} & \frac{\partial \phi_1}{\partial n_2} & \frac{\partial \phi_2}{\partial \theta} \\ \frac{\partial \phi_2}{\partial n_1} & \frac{\partial \phi_2}{\partial n_2} & \frac{\partial \phi_2}{\partial \theta} \\ \frac{\partial \phi_3}{\partial n_1} & \frac{\partial \phi_3}{\partial n_2} & \frac{\partial \phi_3}{\partial \theta} \end{pmatrix} \right|_{\substack{n_1=n_1^* \\ n_2=n_2^* \\ \theta=\theta^*}} \right\}. \tag{20}$$

Checking stability for the monomorphic equilibria we have

$$\det(J(0,K,\theta^*)-\lambda M) = \begin{vmatrix} \frac{1}{\theta^*}(f(\theta^*-\delta)-f(\theta^*))-\lambda & 0 & 0 \\ -\frac{f(\theta^*-\delta)}{\theta^*} & -\frac{f(\theta^*)}{\theta^*}-\lambda & 0 \\ \int_0^{\theta^*} f(t-\delta)dt-\alpha & 0 & f(\theta^*)K \end{vmatrix}$$

$$= \left(\lambda - \frac{1}{\theta^*}(f(\theta^*-\delta)-f(\theta^*))\right)\left(\lambda+\frac{f(\theta^*)}{\theta^*}\right) f(\theta^*)K = 0, \tag{21}$$

$$\implies \lambda_1 = \frac{1}{\theta^*}(f(\theta^*-\delta)-f(\theta^*)) < 0, \quad \lambda_2 = -\frac{f(\theta^*)}{\theta^*} < 0, \tag{22}$$

$$\det(J(K,0,\theta^*)-\lambda M) = \begin{vmatrix} -\frac{f(\theta^*-\delta)}{\theta^*}-\lambda & -\frac{f(\theta^*)}{\theta^*} & 0 \\ 0 & \frac{1}{\theta^*}(f(\theta^*)-f(\theta^*-\delta))-\lambda & 0 \\ 0 & \int_0^{\theta^*} f(t)dt-\alpha & f(\theta^*-\delta)K \end{vmatrix}$$

$$= \left(\lambda+\frac{f(\theta^*-\delta)}{\theta^*}\right)\left(\lambda-\frac{1}{\theta^*}\left(f(\theta^*)-f(\theta^*-\delta)\right)\right) f(\theta^*-\delta)K = 0, \tag{23}$$

$$\implies \lambda_1 = -\frac{f(\theta^*-\delta)}{\theta^*} < 0, \quad \lambda_2 = \frac{1}{\theta^*}(f(\theta^*)-f(\theta^*-\delta)) < 0, \tag{24}$$

$\lambda_1$ and $\lambda_2$ are negative, since $f(\theta^*-\delta) > f(\theta^*)$ at $n_1^* = K$. Therefore, $n_1^* = K$ is stable. Similarly, $f(\theta^*) > f(\theta^*-\delta)$ at $n_2^* = K$, and so it is also stable. Considering the polymorphic equilibrium (where $f(\theta^*-\delta) = f(\theta^*)$), we have

$$\det(J(n_1^*, n_2^*, \theta^*)-\lambda M) = \begin{vmatrix} \frac{f(\theta^*)n_1^*}{\theta^* K}-\lambda & \frac{f(\theta^*)n_1^*}{\theta^* K} & \frac{n_1^* n_2^*}{\theta^* K}(f'(\theta^*-\delta)-f'(\theta^*)) \\ \frac{f(\theta^*)n_2^*}{\theta^* K} & \frac{f(\theta^*)n_2^*}{\theta^* K}-\lambda & \frac{n_1^* n_2^*}{\theta^* K}(f'(\theta^*)-f'(\theta^*-\delta)) \\ \int_0^{\theta^*} f(t-\delta)dt-\alpha & \int_0^{\theta^*} f(t)dt-\alpha & n_1^* f(\theta^*-\delta)+n_2^* f(\theta^*) \end{vmatrix}$$

$$= \left(\frac{f(\theta^*)}{\theta^*}-\lambda\right)\left(\lambda f(\theta^*)K - \frac{n_1^* n_2^*}{\theta^* K}(f'(\theta^*-\delta)-f'(\theta^*))\left(\int_0^{\theta^*} f(t-\delta)-f(t)dt\right)\right) = 0, \tag{25}$$

$$\implies \lambda_1 = \frac{f(\theta^*)}{\theta^*} > 0, \quad \lambda_2 = \frac{n_1^* n_2^*(f'(\theta^*-\delta)-f'(\theta^*))\left(\int_0^{\theta^*} f(t-\delta)-f(t)dt\right)}{\theta^* f(\theta^*)K^2} < 0, \tag{26}$$

since $f'(\theta^* - \delta) - f'(\theta^*) > 0$ and $\int_0^{\theta^*} f(t-\delta)dt - f(t)dt < 0$. Thus, the polymorphic equilibrium is unstable.

##### D.2 Evolutionary invasion analysis

The adaptive dynamics analysis of the Lotka-Volterra DAE gives the same qualitative analysis as the discrete model. Taking the dominant eigenvalue of Eq. 22 and differentiating, we find

$$\frac{\partial}{\partial \delta}\left(\frac{f(\theta^*_{\text{res}} - \delta)}{\theta^*_{\text{res}}}\right)\bigg|_{\delta=0} = -\frac{f'(\theta^*_{\text{res}})}{\theta^*_{\text{res}}}, \quad \frac{\partial}{\partial \delta}\left(\frac{f(\theta^*_{\text{res}} + \delta)}{\theta^*_{\text{res}}}\right)\bigg|_{\delta=0} = \frac{f'(\theta^*_{\text{res}})}{\theta^*_{\text{res}}}, \tag{27}$$

for a delaying and non-delaying mutant, respectively, where $\theta^*_{\text{res}}$ is the equilibrium time for the resident. When $\theta^*_{\text{res}} = \theta^\dagger_{\text{res}}$, the time at which the resident's schedule peaks, we have

$$\frac{\partial^2}{\partial \delta^2}\left(\frac{f(\theta^*_{\text{res}} - \delta)}{\theta^*_{\text{res}}}\right)\bigg|_{\delta=0} = \frac{f''(\theta^\dagger_{\text{res}})}{\theta^\dagger_{\text{res}}} < 0, \quad \frac{\partial^2}{\partial \delta^2}\left(\frac{f(\theta^*_{\text{res}} + \delta)}{\theta^*_{\text{res}}}\right)\bigg|_{\delta=0} = \frac{f''(\theta^\dagger_{\text{res}})}{\theta^\dagger_{\text{res}}} < 0, \tag{28}$$

#### E Further numerical results

##### E.1 Initial conditions: Gaussian random field

To explore the effects of spatial correlation in the initial conditions, $p$ was drawn from an $L \times L$ Gaussian random field normalized to $[0, 1]$ (and $N = 5$ across space). We generated the Gaussian random field by R's gstat package (Pebesma, 2004; Gräler et al., 2016). These results show that initial clustering by types can promote polymorphisms, allowing groups to reach a monomorphic population at the carrying capacity before being invaded by the other type.

Fig. SI-1 depicts the time series of the spatial average of the number of delaying and non-delaying individuals. This figure explains the oscillations we observe in the time series of the proportion of individuals. Once the carrying capacity is reached, the benefit of increased growth has less of an effect and the system is dominated by the between type competition dynamics and diffusion.

Fig. SI-2 depict snapshots at $t = 1000$ for $D = 0.001$. The time series are plotted in Fig. SI-3 for $D = 0.01$ and in Fig. SI-4 for $D = 0.001$.

##### E.2 Initial conditions: U-shaped distribution

Fig. SI-5 depicts the time series of the spatial average of the total number of individuals, $N$. Figs. SI-6 and SI-7 depict snapshots at $t = 1000$ for $D = 0.01$ and $D = 0.001$, respectively. Figs. SI-8 and SI-9 depict the time series until $t = 5000$ for $D = 0.01$ and $D = 0.001$, respectively.

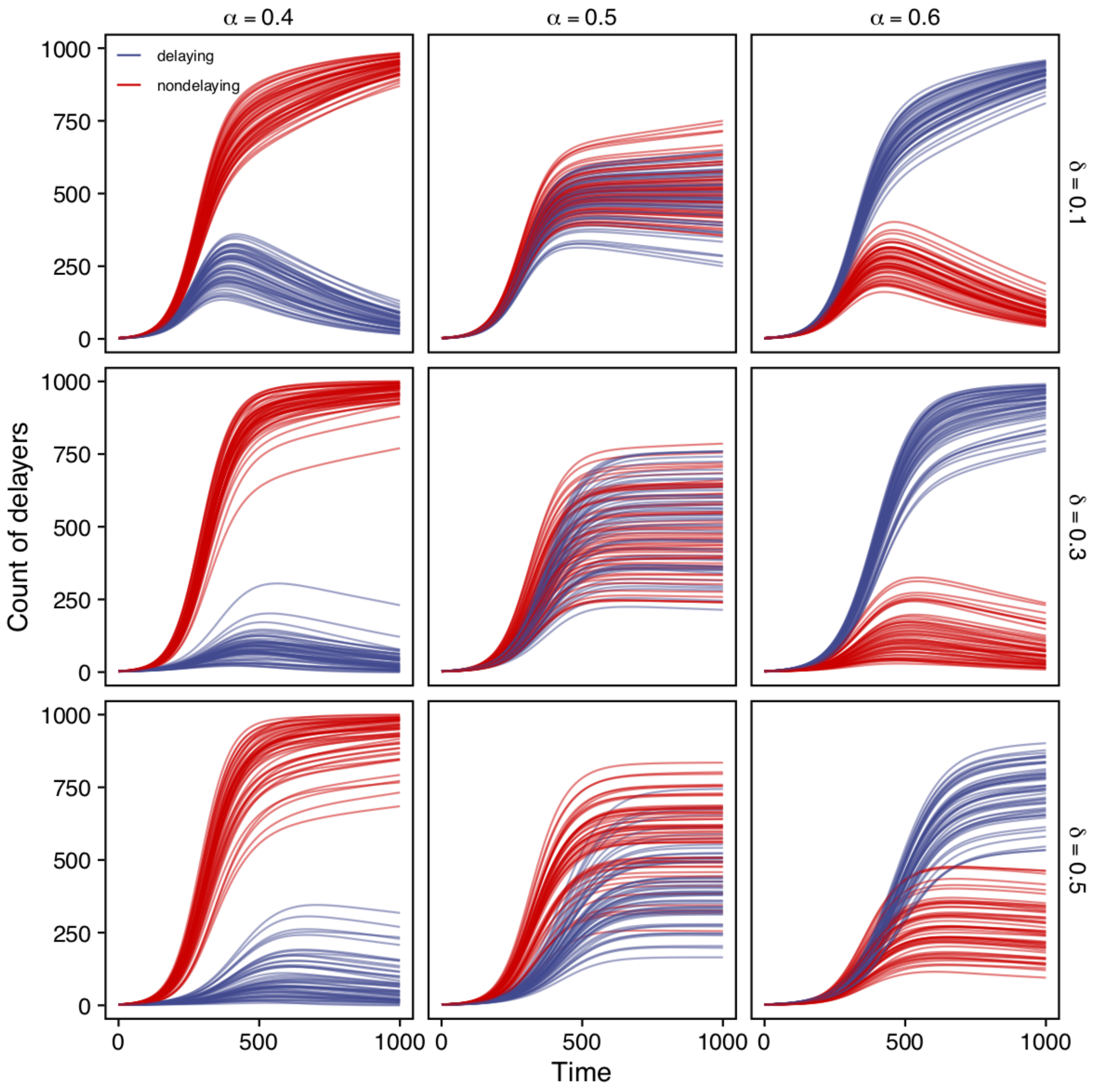


**Figure SI-1.** 50 time series of the spatial average of the number of delaying and non-delaying individuals for low, intermediate, and high $\alpha$ (columns), and where delayers delay contributions with $\delta \in \{10\%, 30\%, 50\%\}$. $D = 0.01$. The initial proportion of delayers were drawn from a normalized Gaussian random field.

James F Crow, Motoo Kimura, et al. An introduction to population genetics theory. *An introduction to population genetics theory.*, 1970.

Benedikt Gräler, Edzer Pebesma, and Gerard Heuvelink. Spatio-temporal interpolation using gstat. *The R Journal*, 8:204–218, 2016. URL `https://journal.r-project.org/archive/2016/RJ-2016-014/index.html`.

Ernst Hairer and Gerhard Wanner. *Solving ordinary differential equations II*. Springer Berlin Heidelberg, 1996.

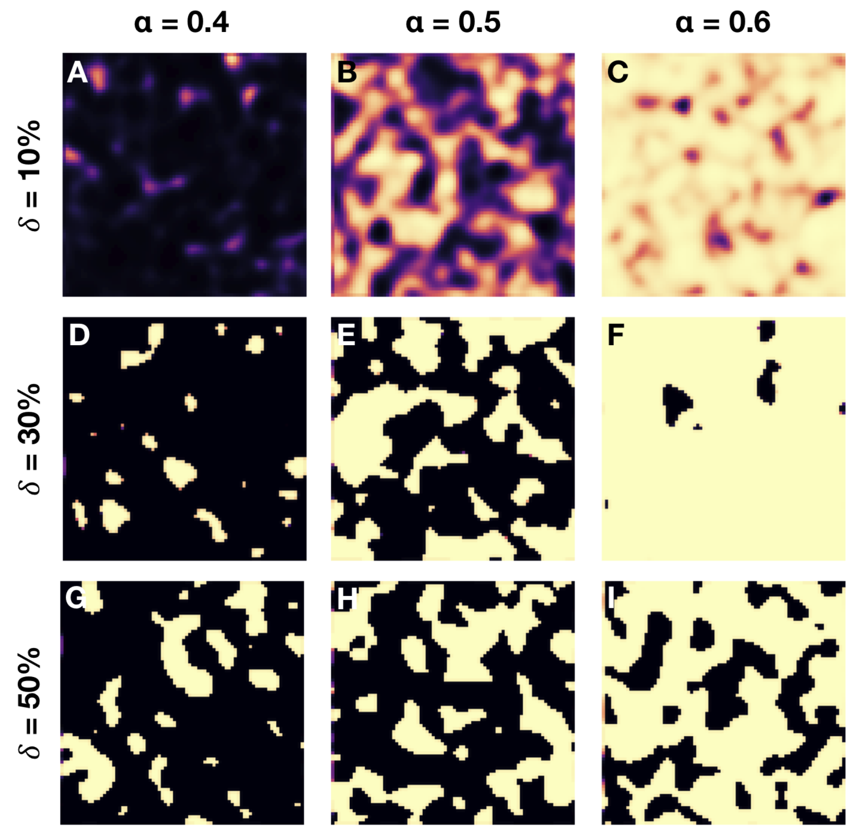


**Figure SI-2. Delayers and non-delayers co-exist for a range of parameters.** Shown are the spatial polymorphisms of delayers (yellow) and non-delayers (black) for the multi-group, multi public goods game model for low, intermediate, and high $\alpha$ (columns), and where delayers delay contributions with $\delta \in \{10\%, 30\%, 50\%\}$. $D = 0.001$. The initial proportion of delayers were drawn from a normalized Gaussian random field. These snapshots are taken at time $t = 1000$.

Edzer J. Pebesma. Multivariable geostatistics in S: the gstat package. *Computers & Geosciences*, 30: 683–691, 2004.

Christopher Rackauckas and Qing Nie. Differentialequations. jl – a performant and feature-rich ecosystem for solving differential equations in julia. *Journal of Open Research Software*, 5(1), 2017. DOI:http://doi.org/10.5334/jors.151.

Ricardo Riaza. A matrix pencil approach to the local stability analysis of non-linear circuits. *International Journal of Circuit Theory and Applications*, 32(1):23–46, 2004.

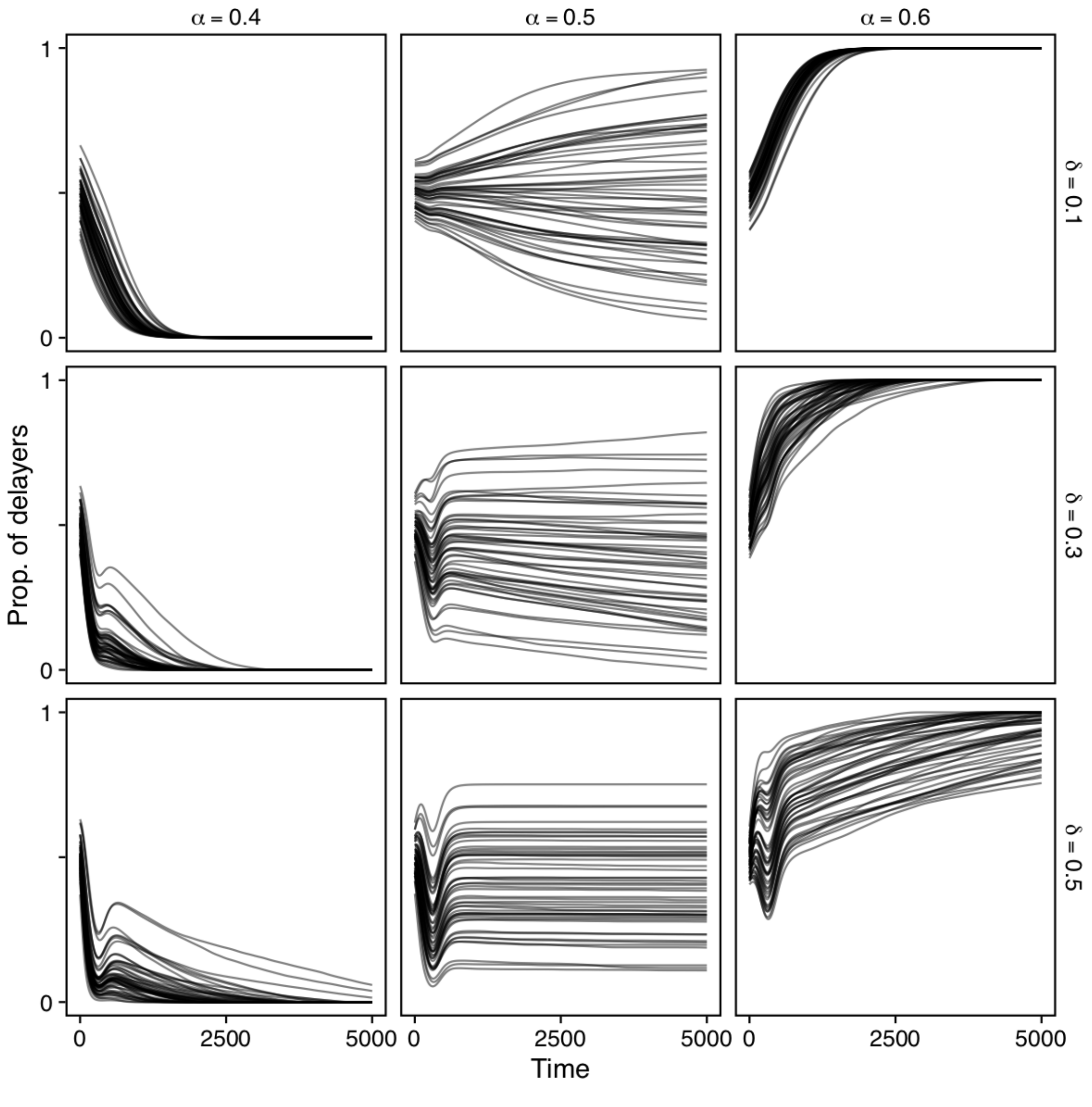


**Figure SI-3.** $50$ time series of the spatial average of $p$ for low, intermediate, and high $\alpha$ (columns), and where delayers delay contributions with $\delta \in \{10\%, 30\%, 50\%\}$. $D = 0.01$. The initial proportion of delayers were drawn from a normalized Gaussian random field.

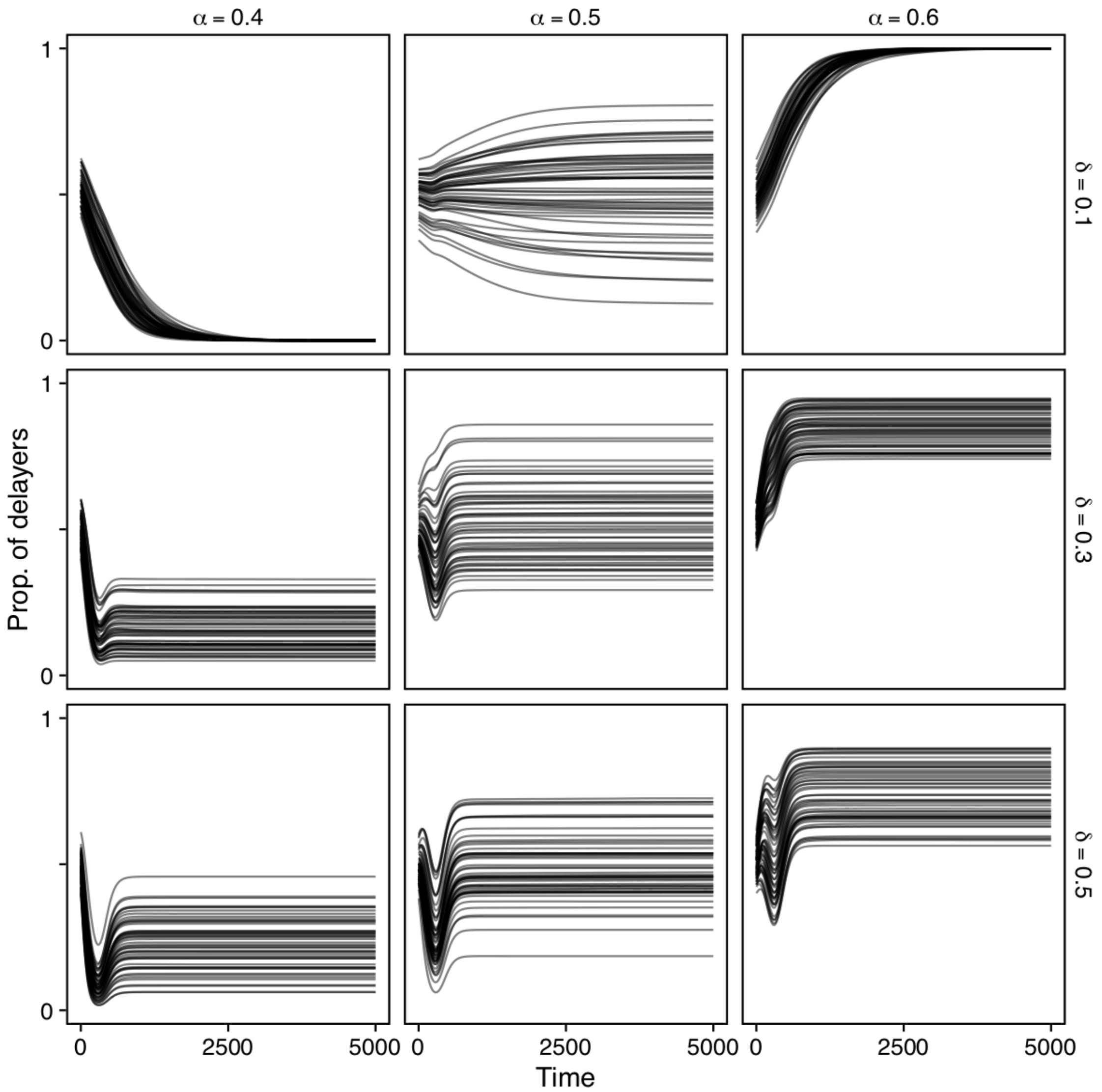


**Figure SI-4.** $50$ time series of the spatial average of $p$ for low, intermediate, and high $\alpha$ (columns), and where delayers delay contributions with $\delta \in \{10\%, 30\%, 50\%\}$. $D = 0.001$. The initial proportion of delayers were drawn from a normalized Gaussian random field.

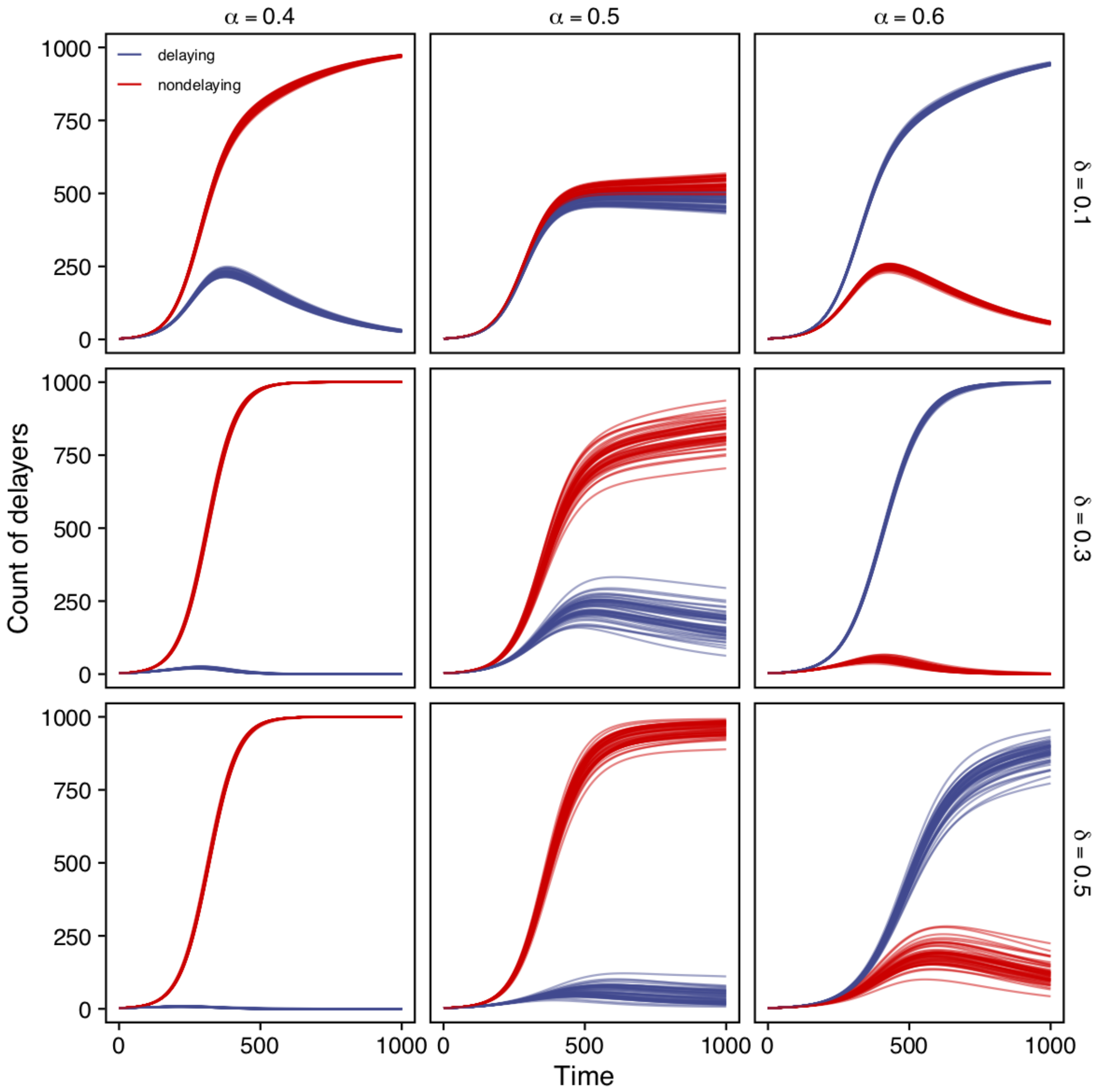


**Figure SI-5.** 50 time series of the spatial average of the number of delaying and non-delaying individuals for low, intermediate, and high $\alpha$ (columns), and where delayers delay contributions with $\delta \in \{10\%, 30\%, 50\%\}$. $D = 0.01$. The initial proportion of delayers were drawn from the Beta distribution with parameters $(0.5, 0.5)$.

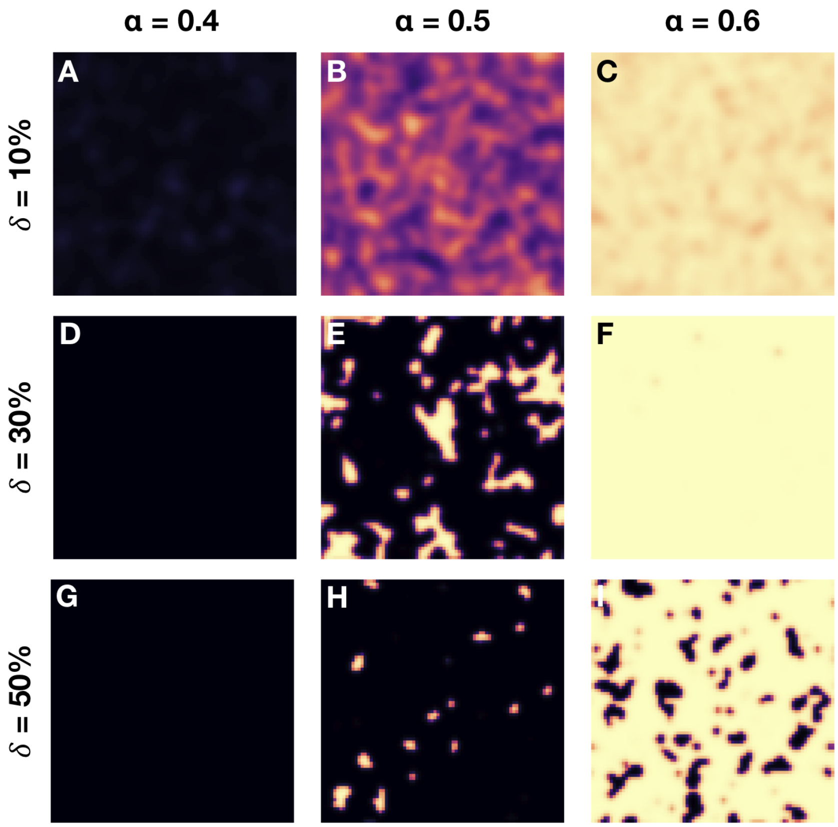


**Figure SI-6. Delayers and non-delayers co-exist for a range of parameters.** Shown are the spatial polymorphisms of delayers (yellow) and non-delayers (black) for the multi-group, multi public goods game model for low, intermediate, and high $\alpha$ (columns), and where delayers delay contributions with $\delta \in \{10\%, 30\%, 50\%\}$. $D = 0.01$. The initial proportion of delayers were drawn from the Beta distribution with parameters $(0.5, 0.5)$. These snapshots are taken at time $t = 1000$.

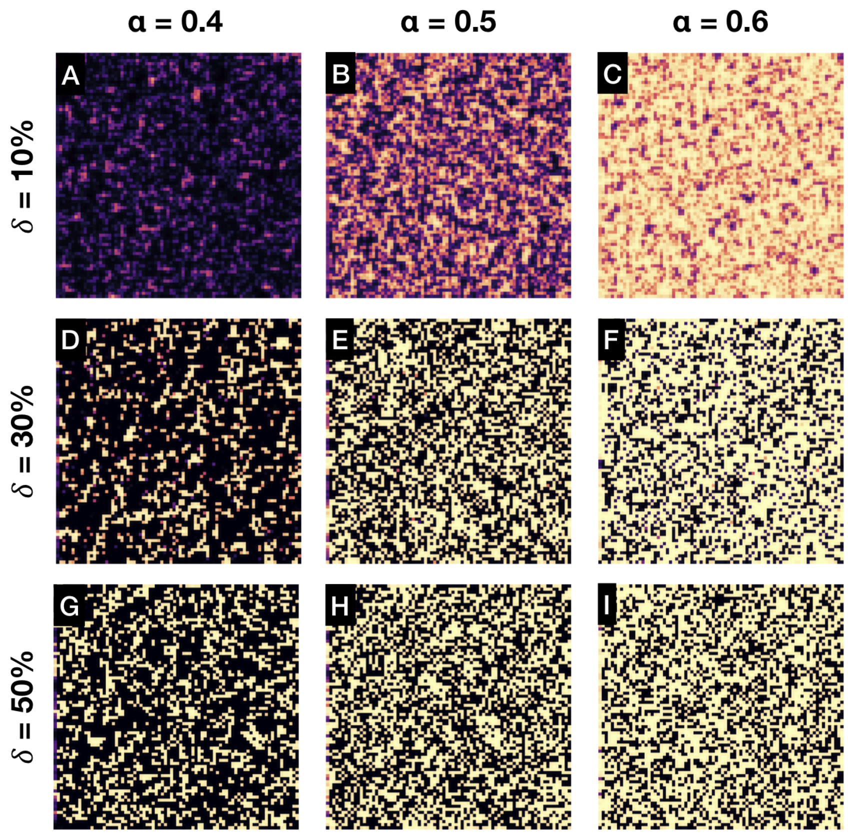


**Figure SI-7. Delayers and non-delayers co-exist for a range of parameters.** Shown are the spatial polymorphisms of delayers (yellow) and non-delayers (black) for the multi-group, multi public goods game model for low, intermediate, and high $\alpha$ (columns), and where delayers delay contributions with $\delta \in \{10\%, 30\%, 50\%\}$. $D = 0.001$. The initial proportion of delayers were drawn from the Beta distribution with parameters $(0.5, 0.5)$. These snapshots are taken at time $t = 1000$.

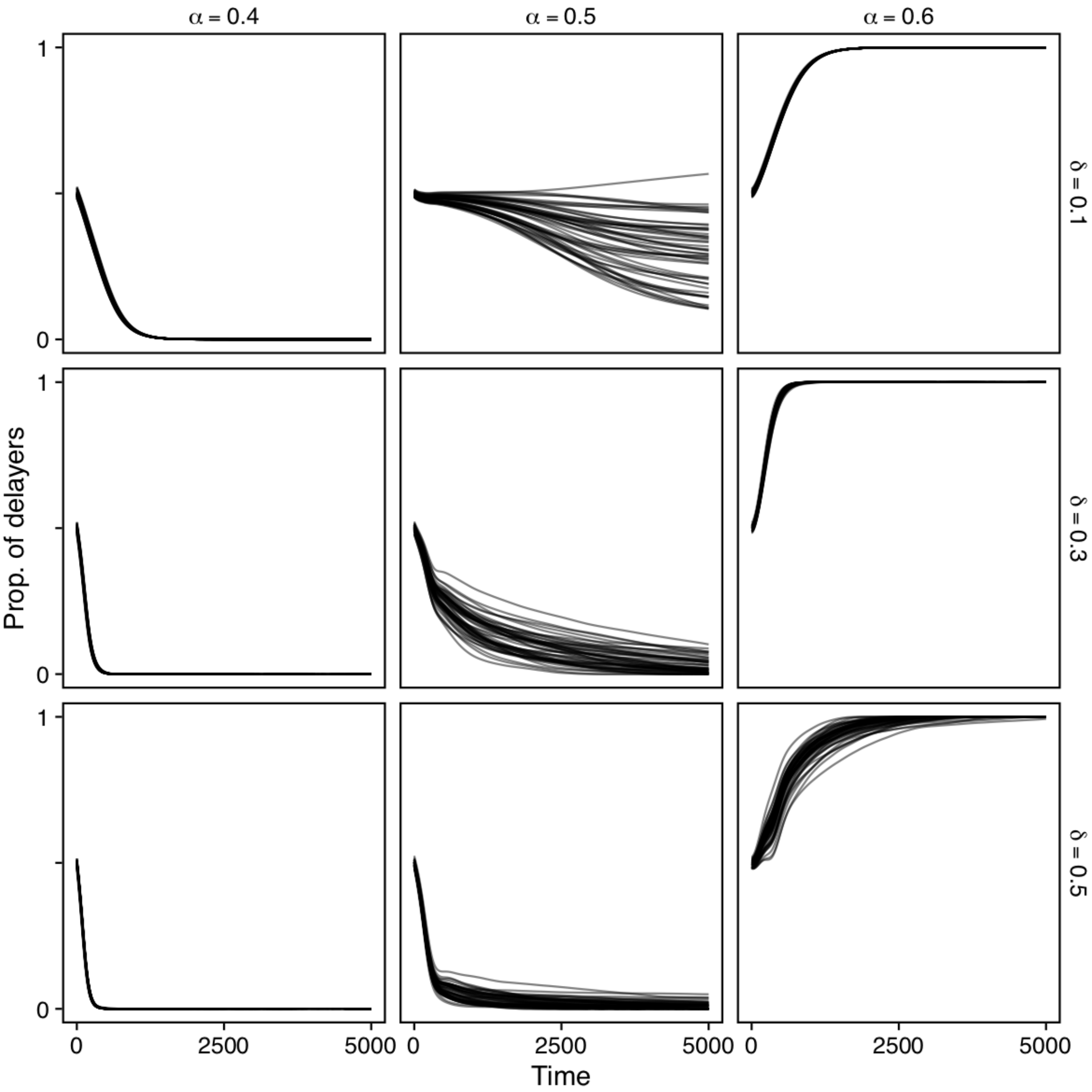


**Figure SI-8.** 50 time series of the spatial average of $p$ (proportion of delayers) for low, intermediate, and high $\alpha$ (columns), and where delayers delay contributions with $\delta \in \{10\%, 30\%, 50\%\}$. $D = 0.01$. The initial proportion of delayers were drawn from the Beta distribution with parameters $(0.5, 0.5)$.

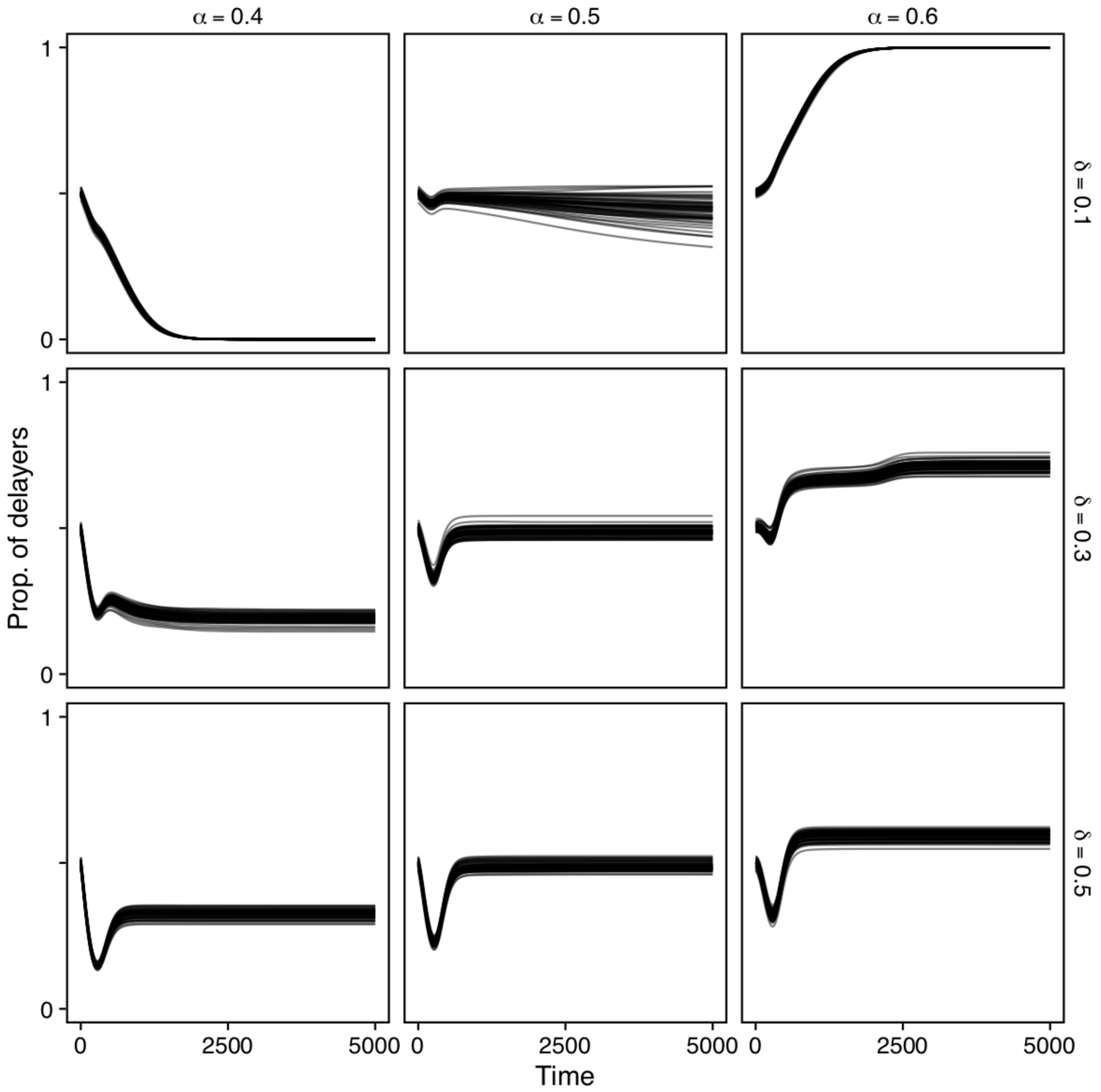


**Figure SI-9.** 50 time series of the spatial average of $p$ (proportion of delayers) for low, intermediate, and high $\alpha$ (columns), and where delayers delay contributions with $\delta \in \{10\%, 30\%, 50\%\}$. $D = 0.001$. The initial proportion of delayers were drawn from the Beta distribution with parameters $(0.5, 0.5)$.
